# Acute effects of subanesthetic ketamine on cerebrovascular hemodynamics in humans: A TD-fNIRS neuroimaging study

**DOI:** 10.1101/2023.01.06.522912

**Authors:** Adelaida Castillo, Julien Dubois, Ryan M. Field, Frank Fishburn, Andrew Gundran, Wilson C. Ho, Sami Jawhar, Julian Kates-Harbeck, Zahra M. Aghajan, Naomi Miller, Katherine L. Perdue, Jake Phillips, Wesley C. Ryan, Mahdi Shafiei, Felix Scholkmann, Moriah Taylor

## Abstract

Quantifying neural activity in natural conditions (i.e. conditions comparable to the standard clinical patient experience) during the administration of psychedelics may further our scientific understanding of the effects and mechanisms of action. This data may facilitate the discovery of novel biomarkers enabling more personalized treatments and improved patient outcomes. In this single-blind, placebo-controlled study with a non-randomized design, we use time-domain functional near-infrared spectroscopy (TD-fNIRS) to measure acute brain dynamics after intramuscular subanesthetic ketamine (0.75 mg/kg) and placebo (saline) administration in healthy participants (*n* = 15, 8 females, 7 males, age 32.4 ± 7.5 years) in a clinical setting. We found that the ketamine administration caused an altered state of consciousness and changes in systemic physiology (e.g. increase in pulse rate and electrodermal activity). Furthermore, ketamine led to a brain-wide reduction in the fractional amplitude of low frequency fluctuations (fALFF), and a decrease in the global brain connectivity of the prefrontal region. Lastly, we provide preliminary evidence that a combination of neural and physiological metrics may serve as predictors of subjective mystical experiences and reductions in depressive symptomatology. Overall, our studies demonstrated the successful application of fNIRS neuroimaging to study the physiological effects of the psychoactive substance ketamine and can be regarded as an important step toward larger scale clinical fNIRS studies that can quantify the impact of psychedelics on the brain in standard clinical settings.

## Introduction

In 2000, the report that ketamine administered intravenously at a subanesthetic dose (0.5 mg/kg) could rapidly alleviate depression symptoms in a handful of treatment resistant patients^1^ was nothing short of a mini-revolution for the treatment of mood disorders. Conventional antidepressants targeting the monoamine system (serotonin, noradrenaline, dopamine) can take weeks to months to act, fail to help as many as 33% of patients^2^, are linked to adverse side effects^3,4^, and it is currently unclear whether they are actually more efficacious in the treatment of depression in adults than a placebo^5^. In contrast to this, the administration of ketamine is a promising treatment option^6,7^. The antidepressant effects of ketamine have been shown to manifest as soon as 40 minutes after administration^8^, which puts it in the category of Rapid Acting Antidepressant Drugs (RAADs). Classic psychedelics have since been added to the RAADs category (e.g. ^9^, for a review see ^10^). As a result, there is now widespread interest in characterizing the physiological and neural effects of psychedelic drugs in general, and ketamine in particular, in a quest to better understand the mechanisms underlying their antidepressant effects^11–13^. Psychedelics are increasingly used for psychedelic-assisted psychotherapy (PAP)^14^, with ketamine-assisted psychotherapy (KAP) as a specific form of PAP^15^.

Ketamine, which has been FDA-approved for medical use since 1970, is increasingly used and researched as a therapeutic approach for depression. Ketamine is a non-competitive N-methyl-D-aspartate receptor antagonist which predominantly affects the glutamatergic system^16^. Ketamine produces strong psychomimetic effects at subanesthetic doses and, therefore, has been used in research as a model for psychosis^17,18^. Ketamine’s delayed effects, in contrast, have been studied in the context of its antidepressant properties^19^. Ketamine and classic psychedelics may exert their antidepressant action through shared mechanisms^20^, despite acting on different neurotransmitter receptors.

Several reviews have been written recently focusing on the action of ketamine^21–24^ and serotonergic psychedelics^20^ at the systems level, as studied with neuroimaging in humans. Many of these reviews highlight the difficulty in establishing a consensus between outcome metrics in extant studies, which typically feature small sample sizes and differences in dosing protocol, control conditions, and analytical methods. Statistical errors that arise from underpowered studies have been discussed extensively, in the context of neuroimaging^25^. Having sufficient data at both the population and individual level is critical when studying psychiatric populations due to the heterogeneity in etiology and in response to different treatments. In the field of depression research, there is burgeoning evidence that alterations within large-scale brain networks are implicated in depressive symptomatology and may thus be used as biomarkers for diagnosis and treatment optimization^26^. However, due to a lack of consistency across studies, such biomarkers have not been established as sufficiently viable in the clinical domain (for a detailed review, see ^27^).

Neuroimaging studies are generally costly and complex. A magnetic resonance imaging (MRI) study must be conducted in the specific location where these expensive devices are installed. This severely limits the ability to run large studies, let alone large and longitudinal studies. However, this is exactly what is needed to make progress in establishing neuroimaging biomarkers of etiology and treatment—recruiting hundreds to thousands of patients and imaging them at several key points in the course of treatment including: at baseline, during the administration of a RAAD, and at several intervals following treatment. While the present study is not aiming to achieve this ideal, it is a proof of concept that such a study is on the horizon.

Recently, Scholkmann & Vollenweider highlighted the “great potential of using optical neuroimaging with functional near-infrared spectroscopy (fNIRS) to further explore the changes in brain activity induced by psychedelics.”^28^. fNIRS neuroimaging is becoming more technically sophisticated and is enjoying increasing popularity and application in the field of neuroscience^29–32^. In the present study we investigated the use of a novel, portable, whole-head time-domain functional near-infrared spectroscopy (TD-fNIRS) system, Kernel Flow1^33^, in a typical outpatient clinical setting, in this case a private practice psychiatry office, to measure changes in cerebrovascular hemodynamics induced by ketamine administration. Besides the scalability of this fNIRS neuroimaging measurement approach due to its ease of use and wearability, it also results in a better “set and setting” for treatment, which has long been considered an important factor in outcome for PAP^9,34^. We note for completeness that a technology such as electroencephalography (EEG)—while not typically considered a neuroimaging technology due to the uncertainty in localizing activity—may also be deployed at scale in clinics, especially devices with active dry electrodes, which are easy to set up and can yield signals of high quality^35^. fNIRS neuroimaging, however, has clear advantages, including the comparability with fMRI measurements^36,37^. After the first fNIRS study with psychedelics was conducted in 2019^38^, this study is the second fNIRS neuroimaging study with psychedelics ever and the first fNIRS neuroimaging study with ketamine ever conducted.

TD-fNIRS systems have been considered the gold standard for optical neuroimaging—they can retrieve more information than the continuous wave (CW-) systems, and in particular they can achieve absolute measurements and higher sensitivity to deeper tissue layers on the human head^39,40^. With these systems, picosecond pulses of light are emitted into the scalp tissue, and arrival times of single photons are measured at nearby detectors. Optical properties of the tissue that lies between a laser source and a detector (which together form a channel) can be inferred across time by analyzing the distribution of the times of flight (DTOFs) of the photons. TD-fNIRS with a high sampling rate enables simultaneous measurement of systemic physiological properties (heart rate and heart rate variability), relative changes in optical properties (which can be linked to hemoglobin oxygenation and concentration changes), and absolute values of optical properties (which can be converted to quantitative hemoglobin oxygenation and concentration measures) under simplifying assumptions.

The current study is a proof of principle of using TD-fNIRS in clinical settings, although this study is not itself a clinical study. We therefore extensively characterized the experience of healthy participants with the system, which they wore while undergoing an intramuscular injection of a saline solution (in their first session), and of a ketamine solution (in their second session). We report on changes in cerebrovascular hemodynamics that we extracted from the recordings: physiological manifestations of ketamine effects on cardiac and respiratory features, absolute hemoglobin concentrations; changes in brain activity as measured by the fractional amplitude of low frequency fluctuations (fALFF)^41^; and changes in global functional connectivity. Finally, we link neural and physiological measurements with participants’ subjective experience as measured by validated surveys.

## Results

### Experimental Design and Data Collection

Participants were fifteen healthy individuals who met eligibility criteria and consented to participation in the study (8 females and 7 males, 24-48 years old; Methods). The study consisted of a single-blind, within-subject experimental design where participants completed four study visits (roughly once a week for four weeks) in a pre-defined order: screening visit, dosing visit 1 (saline, placebo), dosing visit 2 (ketamine), and a follow-up phone call (Fig. 1a; Methods).

**Figure 1.**
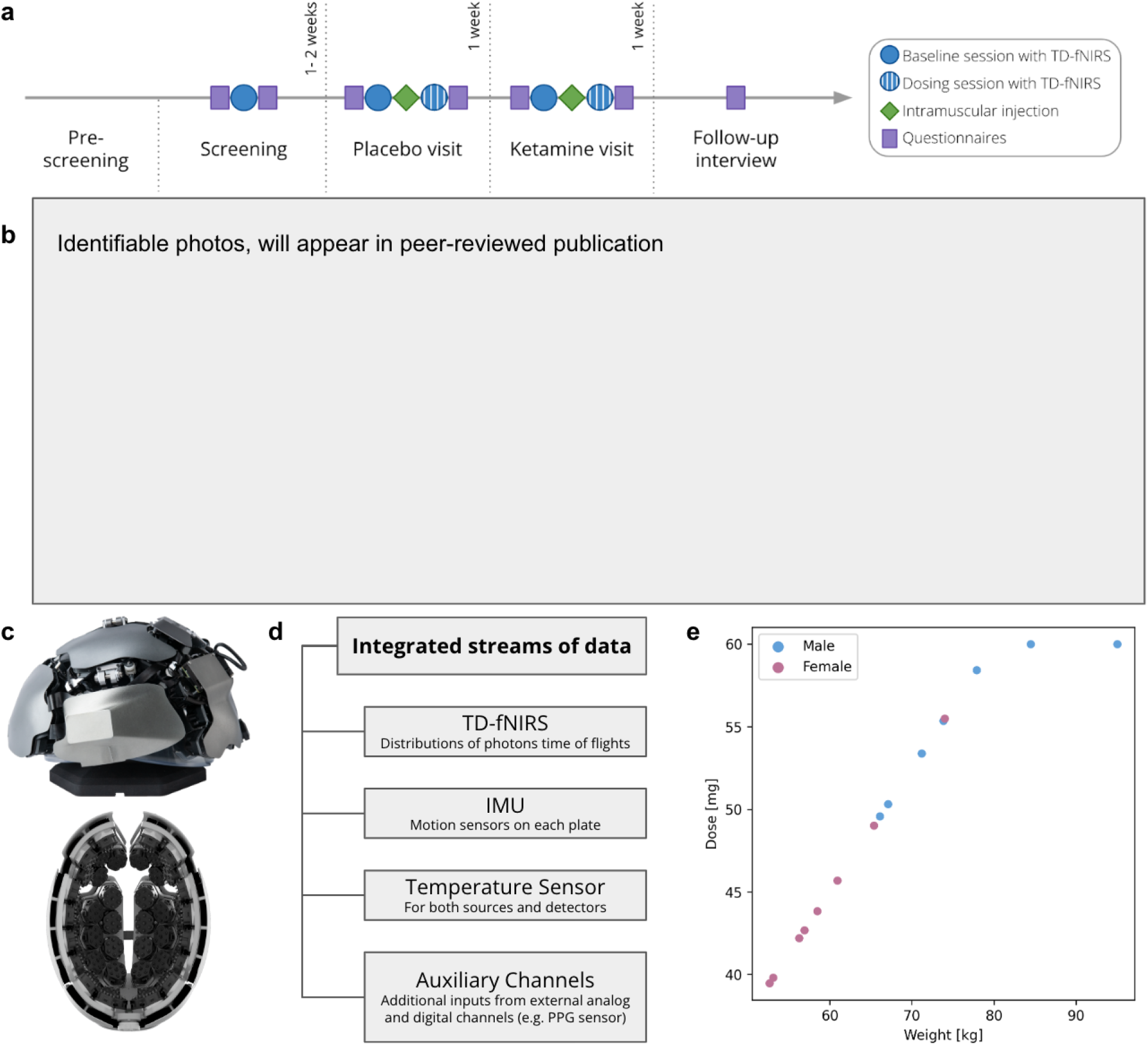
Overview of study design and usage of whole head TD-fNIRS neuroimaging in a clinical environment. **a**. Study Structure. Screening and dosing visits were conducted at the clinical site. The follow up interview was conducted by phone. **b**. Pilot participant wearing the Kernel Flow1 TD-fNIRS system. The TD-fNIRS operator is located to the right outside the image, out of the view of the participant. **c**. The Kernel Flow1 TD-fNIRS headset viewed from the outside (top) and inside (bottom). The headset consists of bilateral frontal, temporal, visual, and sensorimotor plates and a total of 52 modules (each with a dual-wavelength source and six detectors). **d**. Shown are the parallel streams of data that were captured with the Flow1 system in this experiment. **e**. Ketamine dose administered to participants. Each dot represents a participant.

During each dosing visit, participants completed two experimental runs: a baseline run including a short breathing challenge (∼14 minutes), followed by a bolus intramuscular injection to the deltoid of either saline or ketamine (during which participants were partially reclined on a couch for 30 minutes post-dose (Fig. 1b; Methods). Participants wore the Kernel Flow1 TD-fNIRS headset (Fig. 1 c) as well as additional sensors for physiological measurements, including a photoplethysmography sensor (PPG), an electrodermal activity sensor (EDA), and a nasal cannula to measure end-tidal CO2 (EtCO_2_)(Fig. 1d). The amount of administered ketamine was determined by the participants’ weight with a target of 0.75 mg/kg, not to exceed a total dose of 60 mg (reached by two participants) (Fig. 1e)

### Ketamine induces altered states of consciousness and reduces depressive symptomatology

Participants completed surveys at different time points in the course of the entire study (Fig. 1a; Methods; Supplementary Fig. 1), which served to evaluate the subjective (phenomenological) effects of the ketamine vs. saline injection. All participants had higher total scores on the Revised Mystical Experience Questionnaire (RMEQ; paired *T*-test, *p* < 10^−8^), the 5-Dimensional Altered State of Consciousness Rating Scale (5D-ASC; paired *T*-test, *p* < 10^−6^), and the Clinician Administered Dissociative States Scales (CADSS; paired *T*-test, *p*<10^−5^) after ketamine vs. saline. Most participants also had a higher total score on the Brief Psychiatric Rating Scale (BPRS; paired *T*-test, *p* < 10^−3^). In addition, participants completed the Quick Inventory of Depressive Symptomatology (QIDS) questionnaire on three separate occasions, scoring lower one week after ketamine administration, versus one week after saline administration, versus baseline before saline administration (repeated measures ANOVA, *p* < 0.05; Supplementary Fig. 1e).

### Ketamine affects the cardiovascular system, absolute cerebrovascular hemoglobin oxygenation and concentration but preserves respiratory parameters

Often regarded as artifacts, fNIRS data are rich in physiological information from which multiple physiological features can be extracted^42,43^. Although ketamine is thought to preserve baseline respiratory parameters^44^, it is sympathomimetic and has known effects on cardiac signals such as heart rate (HR) and heart rate variability (HRV)^45^. In fact, cardiac measures alone have been suggested as potential biomarkers of treatment outcomes in patients with major depressive disorder (MDD)^45,46^. Consequently, we tested our ability to retrieve several physiological variables from our data and further demonstrate the effects of the active drug as compared to the placebo control.

First, we quantified changes in cardiac signals following ketamine and saline injection. Because this is the first study in which the Kernel Flow1 TD-fNIRS headset was used to derive these signals, we used an auxiliary PPG sensor (attached to the finger) to validate the procedure. Both the PPG sensor and the Flow1 TD-fNIRS system measure pulse rate (PR), which is strongly related to HR^42^. A periodic pulse-like signal (representing the blood volume pulsation) is very apparent when visualizing total photon counts from the 850 nm wavelength of the TD-fNIRS system averaged over well-coupled prefrontal channels, which is aligned with the blood volume pulsation measured by the PPG sensor (Fig. 2a). We found that there was a strong agreement between the instantaneous pulse rate (PR) computed from the PPG and that from the Kernel Flow1 TD-fNIRS system (Fig. 2b; Pearson correlation r=0.93±0.02; mean ± standard error of the mean (SEM) over all sessions) (Methods). We also compared the average PR and pulse rate variation (PRV) extracted from the PPG and the Flow1 system and found that PPG-derived and Flow1-derived metrics were strongly correlated (Fig. 2c). We proceeded to characterize the changes in PR and PRV following ketamine and saline administration. We found a significant increase in PR during the ketamine dosing session (82%) compared to the saline session as well as a significant decrease in PRV (91%) (Fig. 2d)–both findings were confirmed with the external PPG sensor as well and are consistent with those described in prior literature^45^.

**Figure 2:**
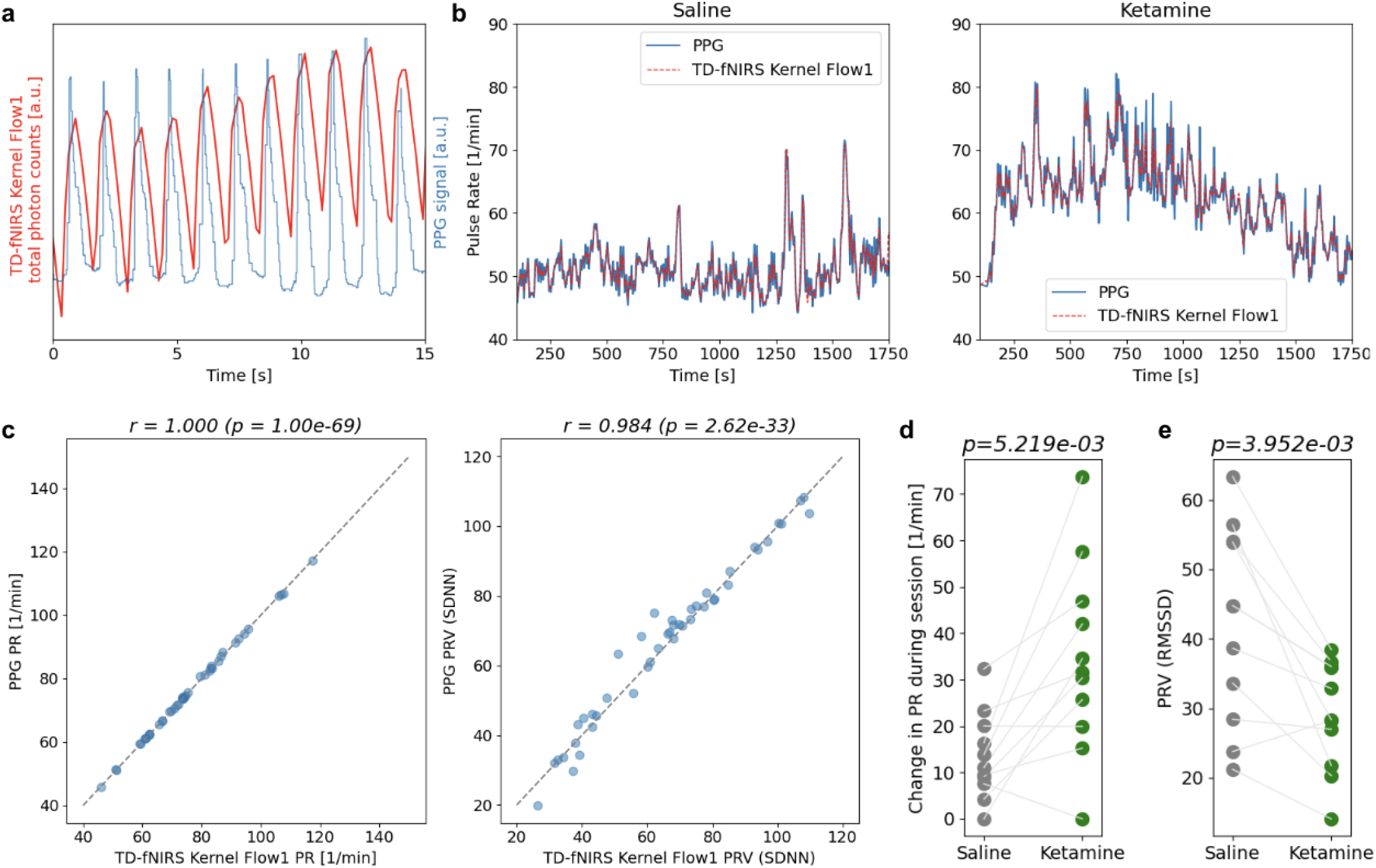
Robust physiological measurements with the Kernel Flow1 TD-fNIRS system reveal significant differences between the ketamine and saline dosing sessions. **a**. The signal from the Kernel Flow1 TD-fNIRS system (total photon counts from 850nm laser averaged over well-coupled prefrontal channels) showed robust heartbeat signals (red) in line with the raw data recorded from an external PPG sensor (blue). Shown is a representative 15 s sample of data. **b**. PR time-series extracted from the TD-fNIRS system (red) and the PPG device (blue) from a representative subject during the saline dosing session (left) and the ketamine dosing session (right). The time-series were smoothed for visualization purposes. **c**. (**Left**) The PR computed from the fNIRS signal was highly correlated with the PPG-based PR (Pearson *r* = 1.00, *p* = 1.01 × 10^−69^; gray dashed line corresponds to the diagonal). (**Right**) Same as Left but for a representative PRV measure (Pearson *r* = 0.98, 2.62 × 10^−33^; SDNN represents the standard deviation of the NN intervals). Note that for this analysis, we made use of all available sessions (both saline/ketamine dosing sessions as well as the baseline runs recorded immediately prior; each dot corresponds to a single recording session). **d**. Maximum change in PR over the course of the dosing session for all participants–the PR increased significantly more during the ketamine than the saline dosing session (paired *T*-test, *p* = 5.22 × 10^−3^). **e**. PRV over the course of the dosing session for all participants–the PRV was significantly lower during the ketamine than the saline dosing session (paired *T*-test, *p* = 3.95 ×10^−3^). Here, PRV is measured by the root mean square of successive differences between normal heartbeats (RMSSD), a common measure of PRV.

Next, we asked whether there were significant differences between ketamine and saline dosing sessions with respect to cerebrovascular hemoglobin. Therefore, we computed the absolute concentrations of oxy- and deoxy-hemoglobin concentrations commonly referred to as HbO and HbR, respectively (Methods). To verify that these signals exhibited expected physiological patterns, we studied the breathing challenge performed during the baseline run prior to each dosing session. Here, a brief hypocapnic state was induced at minute 8 as participants were instructed to breathe rapidly. Indeed, absolute HbO and HbR time-series showed the expected change during the hypocapnic state (Supplementary Fig. 2a). To quantify the change in hemoglobin concentrations as a result of saline or ketamine administration, the session-wise median value of HbO and HbR time-series during the resting-state baseline run were subtracted from those during the dosing sessions. We found a significant increase in HbO (paired *T*-test, *p* = 0.032) and a significant decrease in HbR (paired *T*-test, *p* = 0.021) during the ketamine administration as compared with saline sessions (Supplementary Fig. 2b).

Lastly, we explored other physiological measures that were captured in parallel. Using the capnography time-series recorded from the Nonin capnograph (Supplementary Fig. 3a), we computed the breathing rate during saline and ketamine dosing sessions, and found no significant differences between the two sessions (p>0.1; Supplementary Fig. 3b; Methods). Similarly, no significant differences were observed when comparing the average arterial oxygenation (SpO_2_) and EtCO_2_ values from ketamine sessions to those from saline sessions (Supplementary Fig. 3b), as expected^44^. We did, however, observe an increase in the electrodermal activity (EDA) during ketamine sessions as compared to saline sessions (Supplementary Fig. 3c; saline: 0.19 ± 0.02µS, ketamine: 1.51±0.52µS; mean ± SEM). In addition to the subjective measures of experience following ketamine administration, significant changes in PR, PRV, and EDA provide further corroboration for the effectiveness of the ketamine doses administered in the study.

### Ketamine reduces the whole brain fractional amplitude of low-frequency fluctuations (fALFF)

The fractional amplitude of low-frequency fluctuations (fALFF) in the fMRI-derived blood-oxygen-level-dependent (BOLD) signal has been shown to correlate with cerebral metabolism and is a proxy for local neuronal activity^41^. Abnormal fALFF has been implicated in a variety of conditions, including unipolar and bipolar depression^47–49^. An acute reduction in fALFF following ketamine administration has been reported previously using fMRI^50^. To investigate the changes in fALFF in our study, we first converted the distributions of the times of flight of photons (DTOFs) to the relative concentrations of HbO and HbR (Methods). We then calculated the whole-brain time-varying fALFF for both chromophores using a sliding window approach (duration: 5 min, stride: 10 s; Methods). We found that, at the group level, ketamine dosing sessions showed a marked decrease in fALFF (Fig. 3b), in particular in the later part of the session. Thus, we computed whole-brain fALFF for the first half and second half of the dosing sessions separately. To account for any baseline differences, we used the pre-breathing-challenge portion of the preceding resting-state baseline run to compute a baseline fALFF (Methods). Baseline-normalized whole-brain fALFF was significantly lower in the second half of the ketamine dosing sessions compared to the second half of the saline dosing sessions, for both HbO (paired T-test, *p* = 5.97×10^−3^) and HbR (paired t-test, *p* = 0.025)(Fig. 3c); but not during the first half (*p* > 0.05 for both). The effect of ketamine administration on whole-brain fALFF reaches its maximum around the same time as the PR peaks (at the 500 s mark, on average; Fig. 3a); however, the reduction in fALFF appears to be more sustained than PR, with decreased fALFF persisting for at least 30 min after the injection, at a time when the PR has returned to baseline. This suggests that changes in fALFF and the cardiovascular system are independently affected by the ketamine injection.

**Figure 3:**
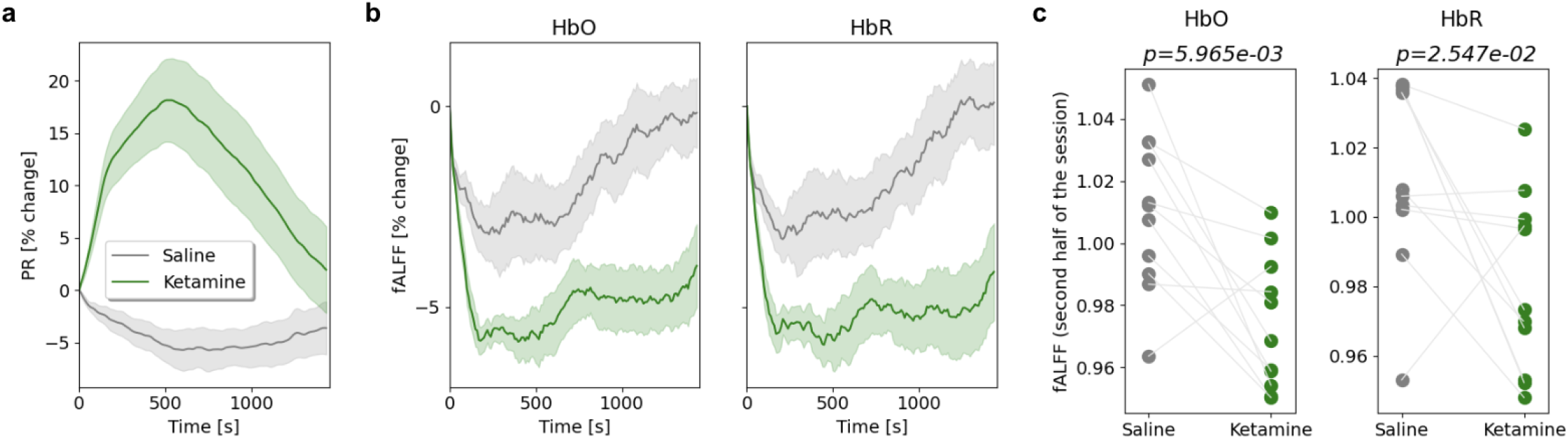
Changes in whole-brain fALFF following ketamine administration. **a, b**. Time-varying PR and fALFF were normalized by the initial value at a given session to obtain the percentage of change throughout the session. Shown are changes in PR (a) and whole-brain fALFF for HbO and HbR (b) during saline (gray) and ketamine (green) dosing sessions (mean ± SEM). **c**. During each saline or ketamine session, whole-brain fALFF (i.e. averaged across the brain) was computed for the first half and second half of the sessions independently and was baseline normalized). Only during the second half of the sessions, fALFF values were significantly lower for the ketamine session compared to the saline session, for both HbO and HbR (paired T-test, *p* = 5.97×10^−3^ and *p* = 0.025 respectively).

### Ketamine acutely reduces global brain connectivity of the prefrontal cortex

The TD-fNIRS system used in this study provides whole head coverage and is thus well-suited to study patterns of cortical connectivity and their acute changes following ketamine administration. Existing fMRI studies focusing on ketamine-induced acute connectivity changes in healthy volunteers found an increase in brain activity^19^: However, this finding is not without controversy. For example, using the measure of global brain connectivity (GBC), some studies report the detection of a global increase in functional connectivity (in particular in prefrontal cortices)^51–54^, but an independent study from another group failed to replicate this finding^55^. Studies using seed-based connectivity report some up- and some down-regulated connections: for example the PCC-mPFC connection is typically found to be downregulated^56,57^; decreased connectivity has also been reported in the visual network^58^; increases and decreases have been reported between the DLPFC and the rest of the brain^59^. In light of the extant literature, our expectations were therefore rather mixed.

To calculate functional connectivity, TD-fNIRS data (i.e. HbO and HbR time-series) underwent further preprocessing (Methods). The group-averaged “dense”, channel-space connectivity matrix (Fig. 4a) exhibited hallmark features of whole-brain functional connectivity such as community structure and homotopic connections. We computed connectivity matrices separately for each session of each visit: baseline and dosing sessions, during the saline and the ketamine visit. For dosing sessions, we focused on a 400 s segment (≈ 6.5 min) at the end of the session (leveraging the observation that fALFF showed the largest differential between ketamine and saline in the later part of the session). We were primarily interested in comparing the ketamine and saline dosing sessions. As with the fALFF analysis, we corrected for any differences in baseline connectivity between the saline and the ketamine dosing sessions by using a segment of the same length (400s) from the immediately preceding baseline session—such baseline differences, whether accidental or related to participants’ expectations, may obscure results or, alternatively, mislead their interpretation.

**Figure 4:**
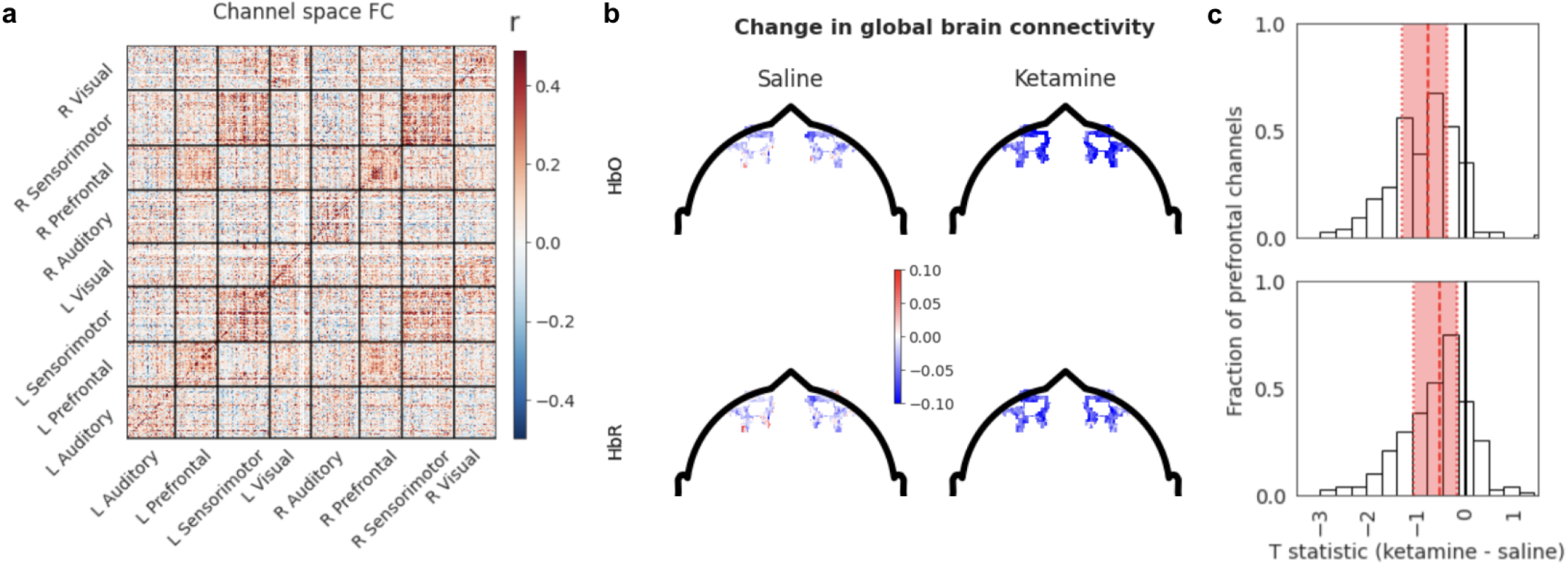
Cortical functional connectivity derived from TD-fNIRS and its acute changes after ketamine vs. saline administration. **a**. Group-level functional connectivity in channel space during the saline dosing session (only HbO is shown). L and R correspond to left and right hemispheres respectively. Sectors correspond to the different plates on the headset (Methods, Fig. 1c). Colorbar indicates the Pearson correlation coefficient used for FC calculations (Methods). **b**. Global brain connectivity (GBC) of the prefrontal channels for saline (left) and ketamine (right) sessions after subtracting baseline GBC for the HbO (top) and HbR (bottom). Note the stronger negative values in the ketamine condition. **c**. The distributions of the group-level *T*-statistic for ketamine vs saline for the HbO (top) and HbR (bottom). The distribution is shifted to the left, indicating a general decrease in GBC. Shown in red are the median and interquartile range.

We started by assessing GBC, a graph-theory inspired metric (a.k.a. weighted degree centrality), which summarizes the connectivity strength of each node of the connectivity matrix^60,61^. We focused our analysis on prefrontal GBC (i.e. quantifying the connectivity of prefrontal channels to the rest of the brain), as the prefrontal area is where GBC has been found to be most affected in previous studies (Methods). As explained above, we baselined GBC for each dosing session using GBC computed during the pre-challenge portion of the resting-state baseline session collected in the same visit. In this setting, we found that ketamine acutely reduces prefrontal GBC compared to saline (Fig. 4b): though no channel or cluster of channels reached significance on its own, the distribution of prefrontal GBC at the group level was significantly shifted to the left (indicating reduction, Fig. 4c). When the participants’ prefrontal GBC was collapsed (by taking the median value of each individual’s distribution over prefrontal channels), indeed, we found a significant reduction in GBC for HbO (*p* = 8.8×10^−3^) and a trend towards reduction for HbR (*p* = 0.05). We further investigated whether any connections were downregulated or upregulated by ketamine, compared to saline: we did not find any connections that were significantly more modulated by ketamine than by saline injection (Methods).

### Physiological and neural features and their relationship with subjective mystical experiences and depressive symptoms

It is argued that one of the predictors of outcome measures in psychedelic therapy is the intensity of mystical experiences reported by the participants^62^. As such, exploring parameters capable of such predictive power can pave the way for treatment optimization. Could a combination of neural and physiological features predict the degree of participants’ mystical experiences? To answer this, we regressed participants’ self-reported RMEQ total scores against time-varying fALFF and PR features (Fig. 5a, Leave One Subject Out cross-validation technique; Methods). The predicted scores on the held-out sessions showed a mean squared error of 1.77 ± 0.44 (mean ± SEM, see Fig. 5b), and the model r-squared on the held-out set was 0.24. Furthermore, we ensured that the performance of the model was above chance level by re-computing the results for 1,000 random permutations of the RMEQ scores (*p* = 0.004; Methods).

**Figure 5:**
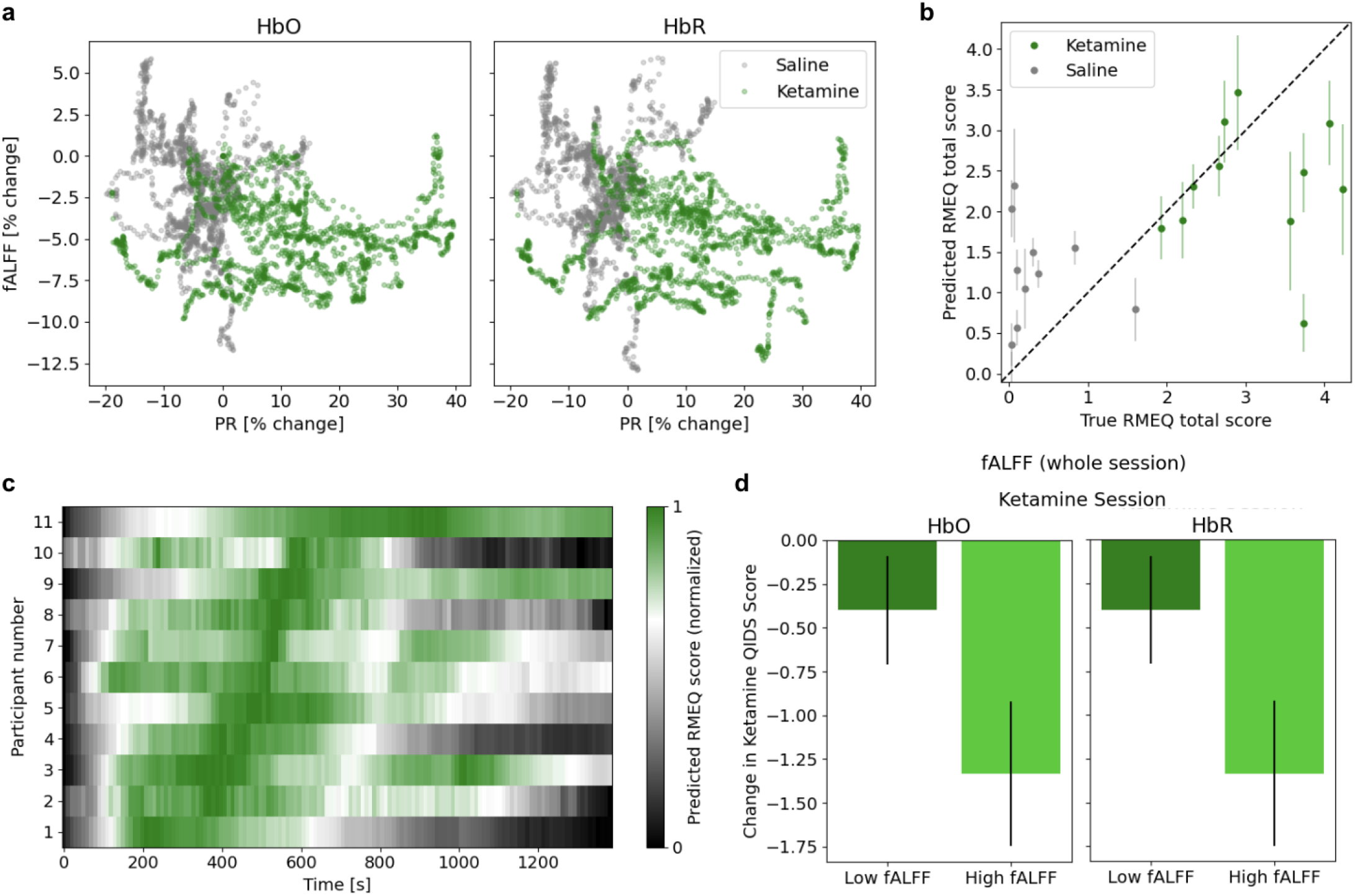
Changes in the magnitude of whole-brain fALFF following ketamine administration. **a**. Time-varying PR and fALFF (normalized by the initial value at a given session) exhibited a separation between ketamine (green cluster) and saline (gray cluster) sessions (note the ketamine cluster in the bottom right corner indicating an increase in the PR and decrease in fALFF). **b**. By using a linear regression model with time-varying PR and fALFF (both HbO and HbR) as input features, we were able to predict the total RMEQ score. Shown are the model predictions on the validation set during each fold (mean ± SEM of the time-varying predictions within each session versus the reported RMEQ total score for the entire session). **c**. We evaluated the trained model on the time-varying features to obtain time-varying RMEQ score predictions (on the validation set). Shown are the normalized prediction scores sorted by the peak time across participants. It appears that different participants may reach their peak of mystical experiences at different time points after ketamine administration. **d**. GEE model revealed that the interaction term between session type (ketamine vs. saline) and whole-brain fALFF (whether it was high or low throughout the session) was a significant factor in the change in QIDS scores (HbO: *p* = 3.53 ×10^−6^, HbR: *p* = 4.76 × 10^−7^). Shown are the change in QIDS scores after the ketamine sessions when we performed a median split on the participants’ fALFF during their ketamine session.

We also calculated the partial correlations between each feature and the RMEQ score while using the other features as covariates, to tease apart the contributions of cerebrovascular and systemic physiological features. We found PR, HbO fALFF, and HbR fALFF to have similar full correlations (0.57, -0.47, -0.43, respectively) as well as partial correlations (0.49, -0.36, -0.32, respectively; *p* < 10^−5^ for all), suggesting that cerebrovascular and systemic physiological signals contain complementary and non-redundant information. When the full model was evaluated at different time points within the ketamine dosing session to obtain time-varying predicted RMEQ scores, it appeared that the peak prediction occurred at different times for each participant (Fig. 5c).

Lastly, we asked whether the change in QIDS scores were related to fALFF during the session. To this end, we modeled the change in QIDS scores as a function of fALFF during dosing sessions (in particular whether session fALFF was high or low, as defined by a median-split), session type (saline/ketamine), and the interaction between the two terms using Generalized Estimating Equations (GEEs). We found that the change in depressive symptoms not only depended on ketamine vs. saline but also on fALFF such that participants with higher fALFF showed a larger reduction in depressive symptoms after ketamine (Fig. 5d; interaction term *p* < 0.001 for both HbO and HbR). Whether the combined changes in PR and fALFF can provide therapeutically relevant biomarkers remains to be explored in future large scale studies in patients.

## Discussion

In the current study, we report on behavioral, systemic physiological, and cerebrovascular measurements from participants who underwent neuroimaging using TD-fNIRS while experiencing altered states of consciousness as a result of subanesthetic ketamine injection in a clinical environment. We compared the acute effects of intramuscular bolus ketamine administration against a placebo (saline administration) condition. We demonstrate reliable physiological measurements of PR and PRV extracted from TD-fNIRS recordings that match those obtained from commercial PPG sensors, thus rendering the use of external sensors to measure cardiac activity unnecessary in future experiments. We also find robust differences between between ketamine and saline conditions: (1) physiological measures: increased PR, decreased PRV, increased absolute HbO and decreased HbR, and elevated EDA (measured by an external sensor) during ketamine dosing, and (2) neural measures: decrease in fALFF and in the global brain functional connectivity of the prefrontal cortex following ketamine administration. These results are generally consistent with findings reported by previous studies (apart from the global functional connectivity result)^45,50,63,64^.

By using a combination of systemic physiological (i.e. PR) and neural (i.e. fALFF) features, we were able to predict participants’ reported RMEQ scores. Further, we found that when the trained model was evaluated throughout the ketamine session, the predicted peak of mystical experience occurred at different times for different individuals. The predicted time-series approximate the expected pharmacokinetics after such an intramuscular bolus and may reflect the time course of the psychoactive effects of ketamine in each participant; to corroborate this, future studies should include continuous subjective ratings of experience and measurements of plasma concentration of the psychoactive substance^65^.

Although we observed a decrease in whole-brain fALFF following ketamine administration, this effect may be region- or network-specific. It has been suggested that fALFF within the default mode network (DMN) is of particular functional importance^48^. Indeed, individuals with MDD show increased fALFF in the left dmPFC, and a ketamine-induced decrease in fALFF suggests a plausible mechanism by which ketamine alleviates depression symptoms^66–68^. Furthermore, we found that fALFF during the ketamine session was a significant factor in the change in depressive symptoms following ketamine administration. It must be borne in mind, however, that while fALFF may be a useful biomarker in predicting treatment outcomes, our study involved only healthy volunteers, and caution must be exercised when extending this measure to the clinical population^67^.

Our result of acutely decreased global functional connectivity in the prefrontal cortex after ketamine administration appears to be at odds with extant literature^53,54^(where a slightly lower dose was used as in our study [0.5 mg/kg vs. 0.75 mg/kg]) though there was a recent report of a failure to replicate past positive findings of a GBC increase in healthy volunteers^55^. In addition, our failure to identify any specific connections with significantly modulated connectivity is at odds with a limited, but certainly mixed^19^, body of reports of acute functional connectivity alterations following ketamine injection. Current studies, this included, have been small and large-scale studies are needed to draw definitive conclusions. We also acknowledge that there are inherent limitations to the whole-head functional connectivity estimates derived with the TD-fNIRS prototype used in this study: for example, not all participants had perfect coupling over the whole head, and the registration of data across participants can be further improved, which may increase statistical power at the group level. Future work is required to ensure improved measurement coverage over the head, and better inter-participant registration capabilities. In this work, reported measures are averaged over large anatomical areas (the whole head or the entire prefrontal cortex), however more sophisticated anatomical modeling and registration methods may allow for more detailed investigations of brain measures. In addition, there are differences in the neuronal-activity-related cerebrovascular changes measured with fMRI and fNIRS. fNIRS is more sensitive to changes in the venous blood fraction and capillaries, compared to the BOLD fMRI signal, and both fMRI and fNIRS measures are a proxy for neural changes^69^. Consequently, systemic physiological activity can have a significant impact on fNIRS signals^70,71^, although our measurement and signal-processing approach was optimized to reduce this impact.

The current study, due to its small sample size and limitations of only measuring cortical connectivity, was not designed to adjudicate conflicts in the existing literature. It was intended as a proof-of-principle for using a novel TD-fNIRS system, in clinical settings. The simplicity of using TD-fNIRS in the clinic, and its ability to measure cortical function (fALFF, functional connectivity) in longitudinal and large-scale studies, may unlock insights on functional brain reorganization, acutely and chronically, following pharmacological interventions. These new insights may be particularly valuable for tailoring KAP or extended to tailoring PAP, more generally.

This study has several further limitations. Our study population was not only small, it was also a sample of healthy volunteers rather than a patient population. If the intent is to understand the mechanism of action of RAADs in clinically depressed patients, a MDD patient sample would be most appropriate. Thus, the results that we report in this work may not apply directly to ketamine treatment of depressed patients. Another limitation is that, while we measured subjective depressive symptomatology one week after ketamine dosing, we did not include brain measurements post-dosing which is when many studies have noted neural changes and psychological effects are measurable (see ^19^ for a review of acute vs. delayed effects of ketamine administration). Our design was also single-blind instead of double-blind; and, because the drug induces such potent subjective experience, most but not all participants were effectively unblinded after participating in the first (saline) dosing session (Supplementary Fig. 1). Such issues with blinding are prevalent amongst studies utilizing full doses of psychedelics, and it appears ketamine is no different. Of course, all these shortcomings are not unheard of in a pre-clinical, proof-of-principle study.

Whole head TD-fNIRS neuroimaging offers an ecologically valid methodology to measure the acute and delayed physiological and neural changes following pharmacological interventions, such as ketamine administration. This scalable technology that captures high-quality cerebrovascular signals and can be deployed widely in clinics may be the solution to the current lack of consistency in neuroimaging investigations of the effects of ketamine administration. But why prefer cerebrovascular measurements to electrophysiological measurements, such as those that can be captured with EEG? While EEG suffers from inherent limitations such as its sensitivity to artifacts (muscle, motion, ambient electromagnetic fields) and lack of spatial specificity, hemodynamic estimates of neural activity are themselves plagued with contamination from systemic physiology^70,71^ and, in the case of fNIRS, limited depth sensitivity. Future studies may benefit from combining multiple imaging modalities (e.g. EEG with TD-fNIRS) to provide a more complete understanding of the impact of psychedelics on the brain in a clinical setting.

## Methods

### Participants and screening procedures

Fifteen healthy participants (8 females and 7 males, age (mean ± SD): 32.4±7.5 years, range 24-48 years) completed four study visits. Participants gave written informed consent before beginning the study in accordance with the ethical review of the Advarra IRB (#Pro00059548), which approved this study. Pre-screening exclusion criteria were history of hepatic or renal disease, history of severe neurological, psychiatric, cardiovascular, or respiratory disease, elevated blood pressure, recent or current substance use disorder (excluding nicotine), suicidal ideation or behavior, or currently taking a central nervous system depressant. Inclusion criteria were prior experience with a classical psychedelic drug (at least one but not more than 20 times), between the ages of 21 and 50, and body mass index between 18-30 kg/m^2^. Participants received monetary compensation.

Once participants consented to participate in the study, they underwent a physical exam, medical history, psychiatric history, and assessment for suicide ideation by the study clinician. Vitals, height, and weight were also measured, and demographic information was collected. Participants then did a short breathing exercise while wearing the Kernel Flow1 TD-fNIRS headset to simulate what future study visits would feel like. Lab tests (complete metabolic panel, pregnancy test, and drug screen) were completed and reviewed by the study clinician no more than 14 days before the first dosing visit. A total of 15 participants completed all of the study visits. Participants were given the option for an additional meeting with the study clinician to ask more detailed questions about what to expect and how to prepare for dosing visits.

### Study design

The study was a single-blind, placebo-controlled, non-randomized design with participants completing study visits roughly once a week for four weeks. The four study visits were always conducted in the same order: a screening visit, two dosing visits, and a follow-up phone call. Dosing visits were always placebo (saline, 0.9% NaCl) first and ketamine second, with the ketamine visit occurring one week (7.1±0.5 days, mean±SD) after the saline visit (Fig. 1a).

Ketamine and saline were administered via bolus intramuscular injection (deltoid muscle). Ketamine dosing was based on participant weight with a target of 0.75 mg/kg, up to the maximum dose of 60 mg. Two participants were administered the maximum dose (Fig. 1e).

### Behavioral and experimental paradigms

At each dosing visit, participants underwent physiological and brain monitoring during a baseline run featuring a short breathing challenge and for 30 minutes after saline or ketamine was injected. Physiological sensors (E4, Empatica, Boston, MA, USA and Lifesense, Nonin, Plymouth, MN, USA) measured respiration rate, SpO_2_, EtCO_2_, PR, and EDA. Brain data was measured with a whole-head TD-fNIRS system^33^ (Flow1, Kernel, Culver City, CA, USA) (for a detailed description see Methods: fNIRS data collection and analyses).

During the baseline run, participants were verbally instructed to breathe normally for the first 8 minutes. Then they were asked to breathe rapidly and deeply until their EtCO_2_ dropped by 10 mmHg (hypocapnic state), which normally took 30 seconds or less. Lastly they were asked to resume breathing normally for 5 minutes. The entire run was approximately 14 minutes.

During the saline and ketamine administration, participants were partially reclined on a couch, supported by pillows. The pilot participant demonstrating the setup in Fig. 1b provided written consent for this photo to be shared in an online open access publication. Participants were given the option to listen to music, which was selected by the clinician and was alyrical as well as melodic (14/15 listened to music during both visits, 1/15 declined music at both visits). Room lights were dimmed. Brain and physiological signals were recorded for 30 minutes post-dose. Vital signs were monitored continuously until 60 minutes post-dose, and then checked at 90 minutes and 120 minutes for safety. Adverse events were recorded as needed throughout the visit. Adverse events occurred in 1 out of 15 (7%) and 9 out of 15 participants (64%) after saline and ketamine injection respectively and included fatigue, high blood pressure, nausea, dizziness, headache, fogginess, thirst, and low SpO2 (Supplementary Fig. 4). No severe adverse events occurred. Participants were allowed to leave with an arranged ride a minimum of 120 minutes post-dose, when deemed safe by the study clinician.

### fNIRS data collection and analyses

#### 1. Data acquisition

Data acquisition was performed using previously described methods^33^. Briefly, the TD-fNIRS system was used to acquire the distributions of the times of flight of photons (DTOFs) from more than 2000 channels across the head. The system consists of 52 modules recording at an effective imaging rate of 7.14 Hz and uses two lasers with wavelengths of 690nm and 850nm. Each module contains a source and six surrounding detectors (at 10mm from the source) and, therefore, multiple channels (a given combination of source-detector) can be formed at various source-detector separations (SDS). The system provides bilateral coverage of prefrontal, parietal, temporal, and occipital regions on separate plates for each region (referred to as: prefrontal, sensorimotor, auditory, and visual respectively) and each hemisphere (Fig. 1c).

In addition to the DTOFs, the Flow1 system is capable of capturing parallel streams of data including: data from 5 inertial measurement units (IMU) distributed throughout the headset as well as temperature sensor data for both sources and detectors that are acquired at all times. Data from auxiliary sources, if connected, will be collected as well (Fig. 1d). For example, in the current study data from an external photoplethysmography (PPG) sensor was acquired and later used for physiological signal validation.

For the fifteen participants who completed all four study visits, technical issues occurred during some sessions leading to partial (3 of 30 dosing sessions) or full (1 of 30 dosing sessions) data loss (Supplementary Fig. 5a).

#### 2. Data preprocessing and feature extraction: relative hemoglobin concentrations

We implemented a channel selection procedure using both a histogram shape criteria as well as the presence of physiological signals^33^. The percentage of retained channels refers to the number of usable channels of neural data and can vary based on the contact between the headset and participants’ head. This percentage is reported for the participants when considering sex, skin type, or hair color and, for the vast majority of the participants, the percentage of retained channels fall between 40-80% (Supplementary Fig. 5b).

Next, histograms from the selected channels were used to compute the moments of the DTOFs–specifically the sum, mean, and variance moments^72,73^. DTOF moments underwent further preprocessing using a motion correction algorithm, Temporal Derivative Distribution Repair (TDDR)^74^. TDDR can leave spiking artifacts in the presence of baseline shifts - we detected these using moving standard deviation, and corrected them using cubic spline interpolation^75^. Finally, we detrended the data using a moving average with a 100-second kernel.

The relative changes in preprocessed moments were then converted to relative changes in absorption coefficients for each wavelength using the sensitivities of the different moments to changes in absorption coefficients (moments Jacobians, i.e. derivative of each moment with respect to a change in absorption coefficient), following the procedure recently described in ^40^. To obtain the sensitivities of the different moments, we used a 2-layer medium with a 12-mm superficial layer. We used a finite element modeling (FEM) forward model from NIRFAST^76,77^, and integrated the moment Jacobians within each layer to compute sensitivities.

Finally, we converted the changes in absorption coefficients at each wavelength to changes in oxyhemoglobin and deoxyhemoglobin concentrations (HbO and HbR, respectively) using the extinction coefficients for the two wavelengths and the modified Beer–Lambert law (mBLL)^78^

#### 3. Extraction of physiological signals

a. Pulse rate, pulse rate variability, and breathing rate extracted from TD-fNIRS data: Within each session, we used the sum moment (total counts) of within-module channels (SDS=10mm) from the prefrontal plates (prior to performing short channel regression). Data from channels with high scalp-coupling index^79^ (SCI>0.85, a minimum of 5 channels) were averaged. We used the python heartpy module^80,81^ to extract physiological signals, for example average and instantaneous pulse rate (PR), and pulse rate variability (PRV). In parallel, we recorded data using an auxiliary PPG sensor and physiological data were also extracted using the heartpy module. The physiological signals computed from the NIRS data were then validated against those measured by the PPG sensor. First, the instantaneous PR from the PPG was interpolated to match the sampling rate of the Flow1-derived PR. For each session, the two PR time courses were compared using a Pearson correlation coefficient. Additionally, the average PR and PRV during a given session were compared against each other and the results are reported in Fig. 2c.
b. Physiological measurements from auxiliary physiological sensors: We applied a fast Fourier transform (FFT) on the time series of CO_2_ waveforms measured during the session to obtain power spectra (Supplementary Fig. 3a). Breathing frequency was obtained by locating the peak of the power spectra during each saline and ketamine dosing sessions (Supplementary Fig. 3b). SpO_2_ and EtCO_2_ signals were continuously recorded throughout the dosing sessions and the average value of each time series was computed during each session as additional respiratory features.

#### 4. Absolute chromophore concentrations

The DTOF is the convolution between the time-resolved temporal point spread function (TPSF) and the Instrument Response Function (IRF). As such, the mean and variance moments of the TPSF can be obtained by subtracting the IRF moments from the DTOF moments. We leveraged calibration data for Flow1 that characterized the IRF for each laser in the system as a function of both temperature and optical power, which allowed us to estimate the IRF throughout the recording session by mapping the instantaneous temperature and power setting to the calibrated IRF value. Absolute moments were then computed by subtracting estimated IRF moments (i.e. mean and variance) from DTOF moments recorded during the experimental sessions.

We then used the analytical expression that converts the mean and variance moments to coefficients of absorption for a semi-infinite homogeneous medium^72,73^. This provides an approximate estimate of absorption coefficients as the semi-infinite assumption is an idealization. These absolute estimates of absorption coefficients can be converted to HbO and HbR concentrations with the mBLL, as described above. To obtain a single time course for HbO and HbR, we computed the median value across well-coupled prefrontal channels with a SDS range of [15, 25] mm.

#### 5. fractional Amplitude of Low Frequency Fluctuations (fALFF)

a. fALFF quantification The time series of data from each channel (for each chromophore) was converted to the frequency domain using an FFT. The ratio of the power in the 0.01–0.08Hz frequency range was calculated relative to the full 0–0.25Hz frequency range. For time-varying fALFF analysis, we used a window size of 5 minutes with a stride of 10 seconds and fALFF was calculated for each window (Fig. 3b, 4a). The fALFF time series was then normalized with respect to the initial value within a given session. For fALFF within the different segments of a session (first half, second half), fALFF was computed over the entire time window of interest (Fig. 3c). The values were then normalized by fALFF computed from a 400s segment of data in the immediately preceding baseline session (selected to end before the breathing challenge, i.e. from 100-500s) to correct for any differences in baseline fALFF between the saline and the ketamine dosing sessions. Whole-brain fALFF was then calculated as the average over all channels.
b. RMEQ total score prediction To predict participants’ total RMEQ scores, we used three features of PR, HbO fALFF, and HbR fALFF (all time-varying features normalized by their initial value in a given session). We used a Leave One subject Out (LOO) cross-validation combined with linear regression. On each fold, both saline and ketamine sessions of a given participant were left out as the validation set and the RMEQ scores were regressed against the data from all other subjects; predictions on the validation set are shown in Fig. 5b. The quality of the fit was further quantified using the mean absolute error on the validation set (i.e. the absolute value of the difference between average predicted RMEQ and true RMEQ). To test whether the performance of the model was above chance level, we used a shuffling procedure. Here, the RMEQ scores (“targets”) were randomly shuffled across sessions so that the samples for each session were assigned a new RMEQ score from a randomly selected session. The entire LOO CV training procedure was then repeated using these shuffled targets, yielding a coefficient of determination and mean MSE. This shuffling procedure was repeated 1,000 times. We then calculated a p-value for the better-than-chance significance of our models’ results by computing the percentage of shuffles which yielded a higher coefficient of determination or lower MSE. To tease apart the contributions of the neural and physiological features, partial correlations were also computed. Here, both fALFF features were used as covariates to PR, while only PR was used as a covariate for each of the fALFF features. As we are interested in exploring whether the neural signal contains additional useful information on top of physiological signals, and since the two fALFF features are highly correlated with each other, including one as a covariate for the other would artificially lower its partial correlation.
c. RMEQ prediction time-series To extract time-varying RMEQ total score predictions, the model (full model, using all features) was evaluated at every time point during the ketamine session. Prediction on the validation set for each participant is shown in Fig. 5c.

#### 6. Functional connectivity

a. Connectivity computation The time series of each channel was further processed by applying a high-pass filter using a bank of discrete cosine basis functions, followed by a low-pass filter using an acausal finite impulse response (FIR) filter. The pass band was 0.01-0.1 Hz, as is customary in the functional connectivity literature. Functional connectivity was computed for all pairs of channels as the Pearson correlation coefficient between their time courses, for each chromophore. For the baseline sessions, we selected a 400 s segment of data ending before the breathing challenge, from 100-500 s, to compute connectivity. This segment of data can be considered to be a resting session. For the dosing sessions, we selected a 400 s segment of data, from 1100-1500 s, to compute connectivity. We chose this segment to be near the end of the session (maximal expected neural effect), and of the same duration as the baseline segment.
b. Global brain connectivity For each channel in the prefrontal plates, a connectivity strength score was obtained by averaging all positive connections to other channels measured over the whole head. A connectivity strength map (limited to the prefrontal plates) was thus computed for each session.
c. Statistical comparisons We used the baseline session as a baseline for all connectivity analyses. We reasoned that the metric of interest was a change in the state of participants, and using a within-visit reference allowed us to focus on this change. For functional connectivity comparisons, we computed the median of the within-network connections and between-network connections separately for each network and each participant, then used a *T*-test to assess significance at the group level. For global functional connectivity comparisons, we computed the spatial maps of prefrontal GBC change for the ketamine and the saline sessions for each subject, then used a paired t-test to highlight consistent modulations at the group level. We further looked at the distribution of the GBC changes in the group map (collapsing the spatial dimension) and tested whether the distribution was centered on 0 using one sample *T*-test.

### Survey data collection

Participants completed the Columbia Suicide Severity Rating Scale (CSSRS)^82^ at the beginning and end of each dosing visit for safety reasons. They also completed the Quick Inventory of Depressive Symptomatology (QIDS-SR)^83^ to report depressive symptoms experienced over the previous week at the beginning of each dosing session and during the follow-up phone call. The clinician administered the Brief Psychiatric Rating Scale (BPRS)^84^ and Clinician-Administered Dissociative States Scale (CADSS)^85^ approximately 60 minutes after the placebo or ketamine administration. Participants then completed the Revised Mystical Experience Questionnaire (RMEQ-30)^86^ and the 5-Dimensional Altered States of Consciousness Rating Scale (5D-ASC)^87^. Participants were also asked about their overall experience with wearing the TD-fNIRS headset using a customized survey (Supplementary Fig. 1f).

## Acknowledgments

This study was funded by Cybin.

## Author contributions

Conceptualization: KLP, WCR, RMF, FS. Methodology/Software: FF, MS, SJ, JD, ZMA, JKH, WCH, NM, JP. Formal analysis: ZMA, JD, MT, FF. Investigation: MT, WCR, WCH, JP, AG. Data Curation: MT. Writing - Original Draft: ZMA, JD, KLP, MT. Writing - Review & Editing: All authors. Project administration: MT, AC. Author order was determined alphabetically. Dr. Katherine Perdue agrees to be accountable for all aspects of the work, ensuring that questions related to the accuracy or integrity of any part of the work are appropriately investigated and resolved.

## Data availability statement

The datasets used and/or analyzed during the current study are available from the corresponding author on reasonable request.

