## Supplemental Info for "Acute effects of subanesthetic ketamine on cerebrovascular hemodynamics in humans: A TD-fNIRS neuroimaging study"

### Supplementary Material

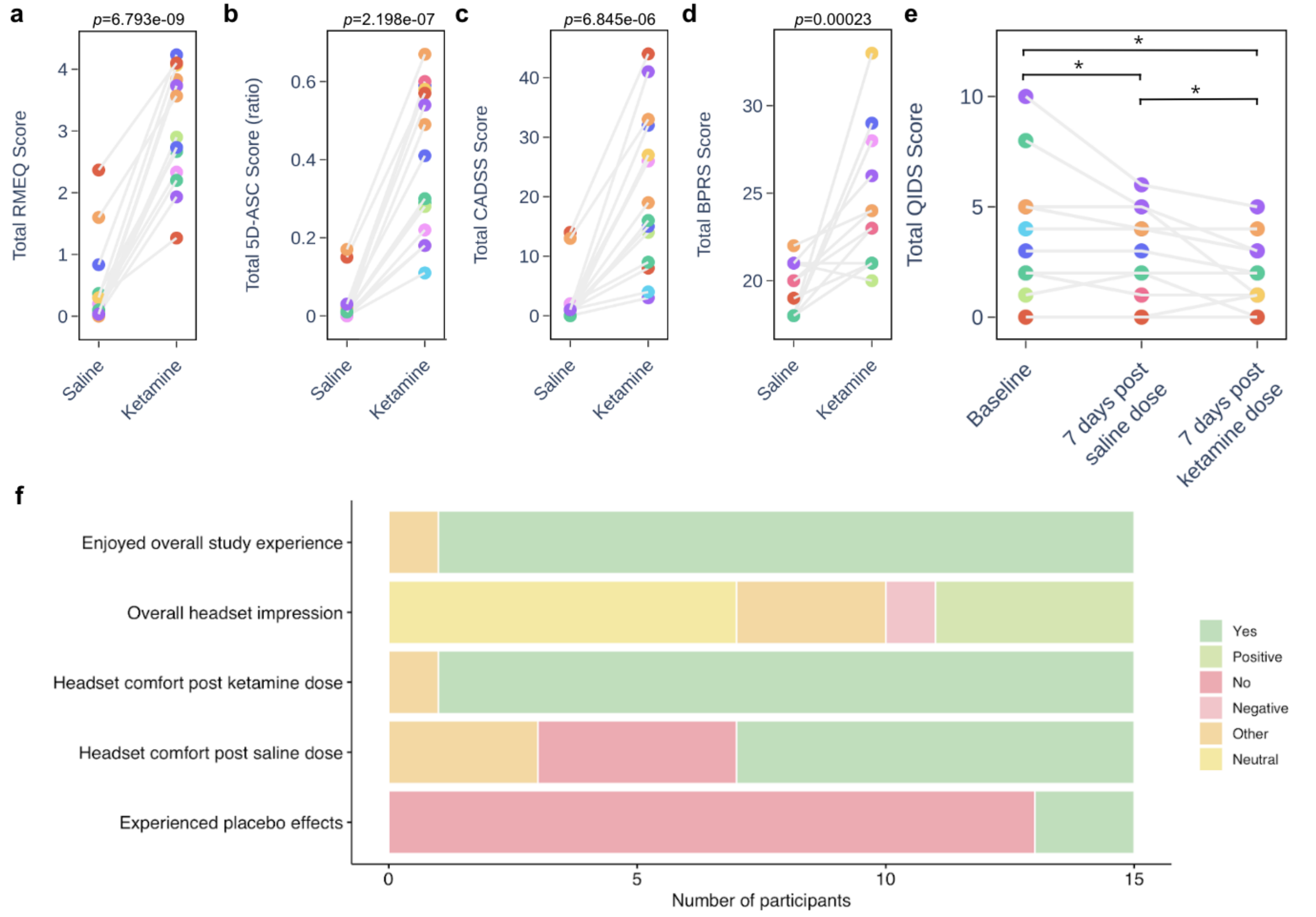


**Supplementary Figure 1: Survey data analysis.**

**a.** Participants' mystical experiences were significantly increased after the ketamine dosing session compared to the saline dosing session. Shown are the RMEQ survey scores, namely the total score that is a composite measure of mystical experiences, positive mood, transcendence, and ineffability (paired t-test p-value$<10^{-8}$). Each dot represents a participant. **b.** Participants' altered states of consciousness were significantly increased after the ketamine dosing session compared to the saline dosing session. Shown are the 5D-ASC survey scores, namely the ratio score that is a composite measure of oceanic boundlessness, dread of ego dissolution, visionary restructuralization, auditory alterations, and vigilance reduction (paired t-test p-value$<10^{-6}$). **c.** A clinician administered survey, CADSS, exhibited significant changes after ketamine administration (paired t-test $6.8\times10^{-6}$). In panels a-c, it is apparent that two study participants experienced a placebo effect. **d.** BPRS, another clinician administered survey, also showed participants' significant changes in psychiatric state after ketamine administration (paired t-test $p=2.3\times10^{-4}$). **e.** Total QIDS scores were measured at different time points and the results from repeated measures ANOVA are shown (* indicates *p-value* < *0.05*). It is worth noting that the majority of the participants (13 out of 15) had few depressive symptoms (scores < 5) and only two of them had mild symptoms (scores between 5 and 10) at the time of the initial visit. **f.** Participants' overall study experience and impression of wearing the headset throughout the study was captured using a customized survey.

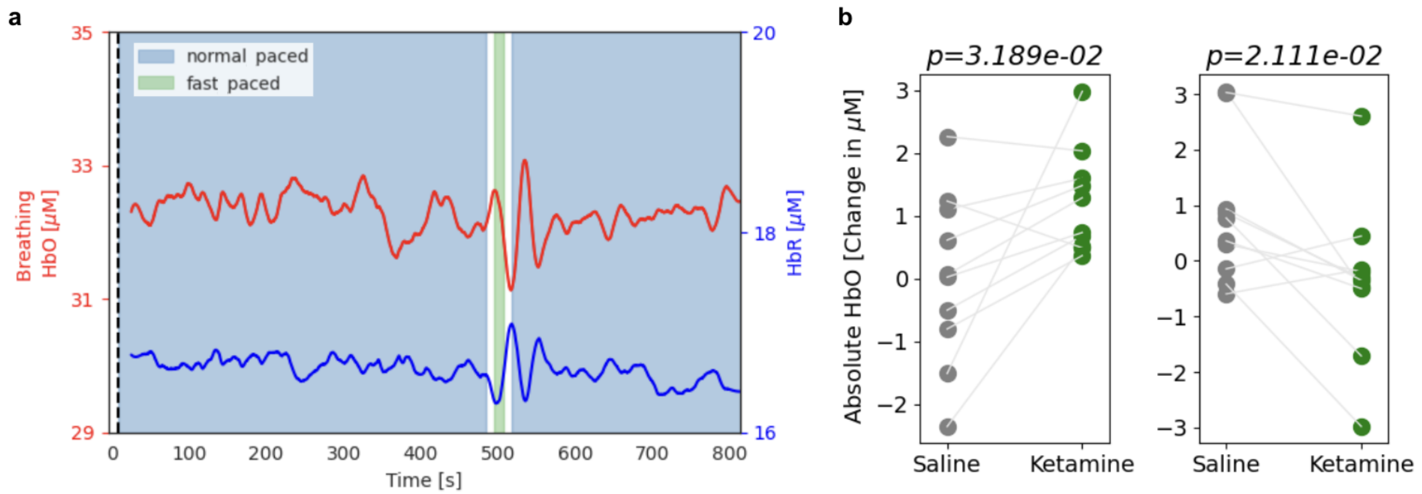


**Supplementary Figure 2: Quantification of changes in absolute oxy- and deoxy-hemoglobin concentration levels. a.** Time-series of HbO (red) and HbR (blue) from a representative breathing experiment. Note the visible change around the time of hypocapnic state (vertical green box). **b.** The median values of HbO and HbR time courses were computed during each session. The value from the resting-state phase prior to each dosing session was used as a baseline and subtracted from that of the dosing session. A significant increase in HbO (left) and a significant decrease in HbR (right) were observed during ketamine compared to saline sessions (shown are paired t-test *p*-values).

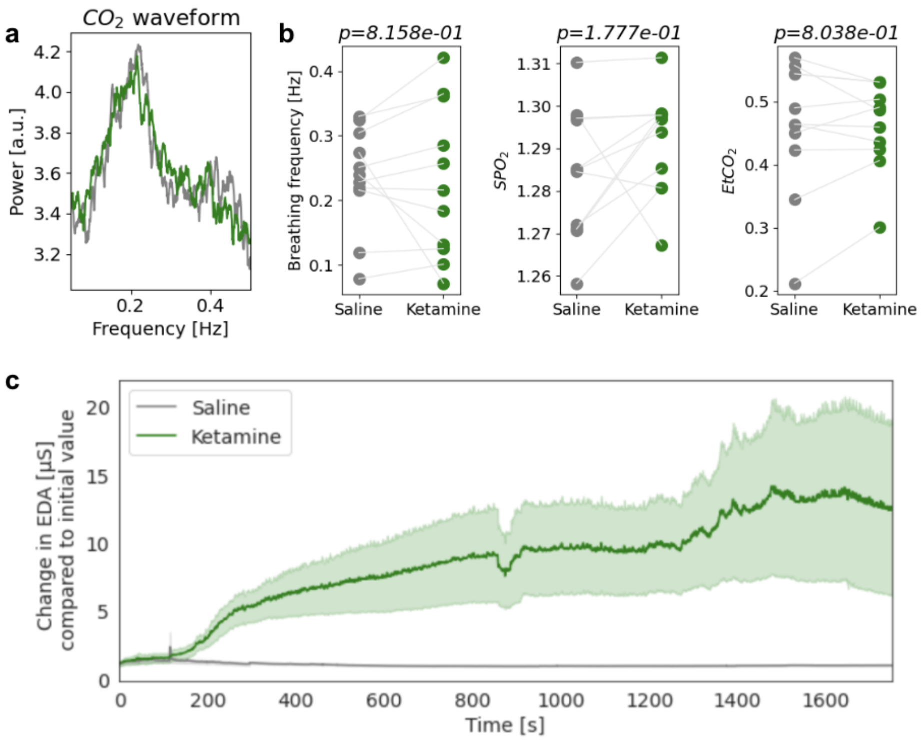


**Supplementary Figure 3: Extended physiological changes between saline and ketamine dosing sessions.**
**a.** Representative power spectrum of the $CO_2$ waveform recorded with the Nonin capnograph. The saline (gray) and ketamine (green) sessions spectra are very similar in the example participant shown. **b.** No significant difference was found between the saline and ketamine dosing sessions in terms of breathing frequency (derived from Nonin data; left), peripheral capillary oxygen saturation ($SpO_2$, middle) or end-tidal $CO_2$ ($EtCO_2$, right); all $p > 0.1$. **c.** Electrodermal activity (EDA) during the dosing sessions (when normalized by the initial value) showed a significant increase during ketamine versus saline sessions (shown are mean ± SEM).

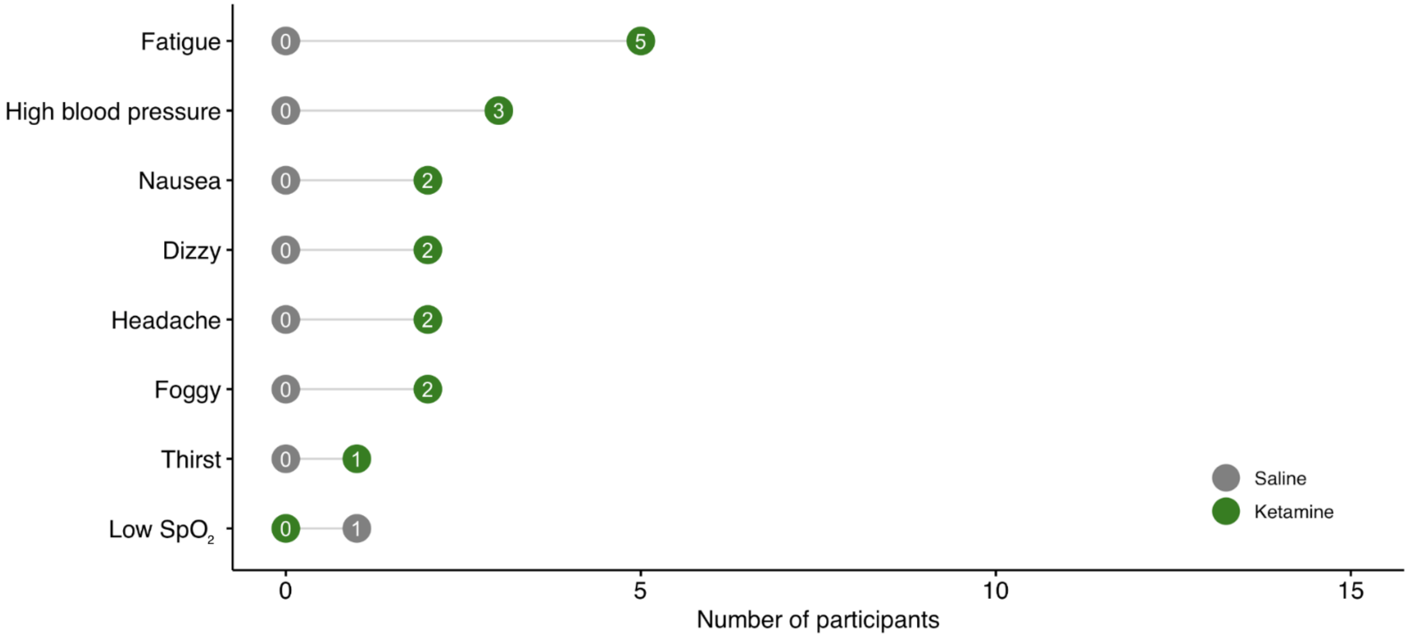


**Supplementary Figure 4: Adverse events.**
Shown are the number of participants who reported having had adverse events after saline (gray) and ketamine (green) administration.

**a**

| | Survey | Physiological | Neural | | |
|---|---|---|---|---|---|
| **Visit** | Complete | Complete | Complete | Partial | Missing |
| Saline | 15 | 15 | 11 | 3 | 1 |
| Ketamine | 15 | 15 | 15 | 0 | 0 |

**b**

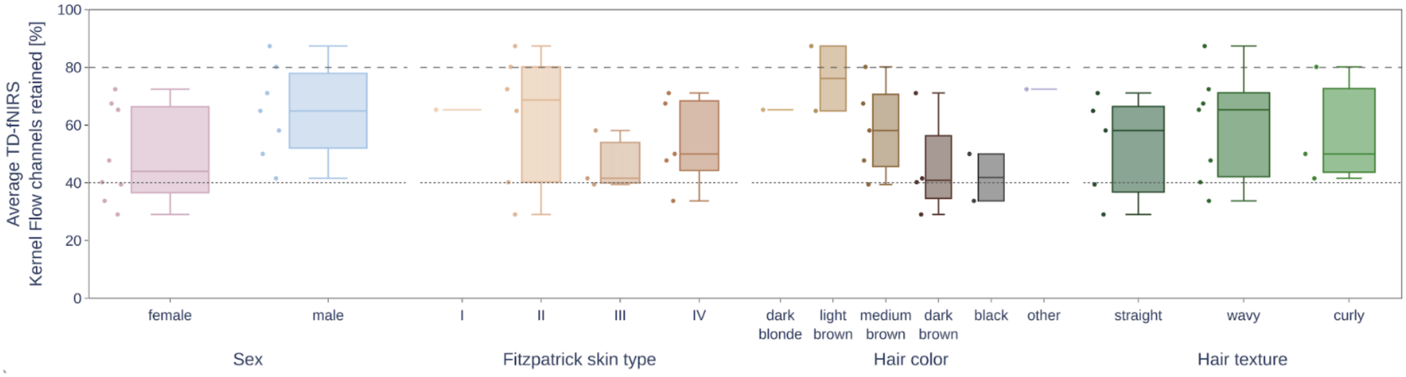


**Supplementary Figure 5: Data quality and completeness.**
**a.** Number of dosing sessions that incurred partial or full data loss due to technical issues. **b.** For all participant demographics, the percentage of retained channels with good coupling (a metric of coverage of measured signals) are shown. The dotted line at 40% and dashed line at 80% indicate quality control thresholds for “good” and “excellent” fractions of retained channels, respectively.
